# *Wolbachia* infection negatively impacts *Drosophila simulans* heat tolerance in a strain- and trait-specific manner

**DOI:** 10.1101/2023.12.18.572256

**Authors:** Liam Ferguson, Perran A. Ross, Belinda van Heerwaarden

**Affiliations:** School of BioSciences, Bio21 Molecular Science and Biotechnology Institute, The University of Melbourne, Parkville, Australia; Section for Bioscience and Engineering, Department of Chemistry and Bioscience, Aalborg University, Aalborg, Denmark

**Keywords:** *Wolbachia*, *Drosophila simulans*, climate change, heat tolerance, CTmax, fertility thermal limits, upper thermal limits

## Abstract

The susceptibility of insects to rising temperatures has largely been measured by their ability to survive thermal extremes. However, until recently, the capacity for maternally inherited endosymbionts to influence insect heat tolerance has been overlooked. Further, the impact of heat on traits like fertility, which can decline at temperatures below the lethal thermal limit has largely been ignored. Here, we assess the impact of three *Wolbachia* strains (*w*Ri, *w*Au, and *w*No) on the survival and fertility of *Drosophila simulans* exposed to heat stress during development or as adults. The impact of *Wolbachia* infection on heat tolerance was generally small and trait/strain specific. Only the *w*No infection significantly reduced survival and fertility of adult males after a heat shock. When exposed to a fluctuating heat stress during development, the *w*Ri and *w*Au strains reduced egg-to-adult survival but only the *w*No infection reduced male fertility. *Wolbachia* densities of all three strains decreased under developmental heat stress, but reductions occurred at temperatures above those that reduced fertility of the host. These findings reveal the complexity of endosymbiont-host-environment interactions and emphasise the necessity to account for endosymbionts and their effect on both survival and fertility when investigating the vulnerability of insects to climate change.

## Introduction

Insects are one of the most economically and ecologically important groups of organisms, responsible for crop pollination, wildlife nutrition, and pest control, among other roles (Losey and Vaughan, 2006). Many species are at risk of extinction due to climate change, which poses a significant threat to food security and ecosystem stability (Hallmann *et al*., 2017; Sánchez-Bayo and Wyckhuys, 2019). Insects and other ectotherms rely on environmental temperature for optimum physiological function, so their global distribution is generally defined by thermal limits (Cossins and Bowler, 1987). Rising temperatures caused by climate change have driven shifts in insect distributions, particularly away from the equator where tropical species are already living close to their upper thermal limit (Parmesan and Yohe, 2003; Deutsch *et al*., 2008; Lenoir and Svenning, 2015). To adapt to rising temperatures, insects can increase their upper thermal limits through short term phenotypic changes (plasticity) or long-term evolutionary adaptation (Hoffmann and Sgró, 2011; Diamond and Martin, 2016; Hoffmann *et al*., 2023). However, evolutionary and plastic responses in insects are often small, limiting their capacity to counter current rates of climate change (Hoffmann *et al*., 2013; Gunderson and Stillman, 2015; Hangartner and Hoffmann, 2016; Kellermann and van Heerwaarden, 2019; Weaving *et al*., 2022).

Until recently, research has focused on insect heat tolerance without considering the potential impact of endosymbionts, which may be powerful sources of additional phenotypic variation (Wernegreen, 2013; Corbin *et al*., 2017; Dunn, 2017; Hector *et al*., 2022). Endosymbionts are intracellular bacteria that are carried by many insect species and are maternally inherited (Weinert *et al*., 2015). Endosymbionts have diverse effects on their host, including pathogen protection, nutrient supplementation, and changes in reproduction (Werren *et al*., 2008; Eleftherianos *et al*., 2013; Newton and Rice, 2020). Among these effects, a growing body of evidence suggests that endosymbionts can alter the heat tolerance of their host, for better or for worse (Russell and Moran, 2006; Brumin *et al*., 2011; Heyworth and Ferrari, 2015; Tougeron and Iltis, 2022). Models predicting the vulnerability of insects to climate change that do not consider endosymbionts may provide inaccurate estimates of insect susceptibility to rising temperatures.

Insect heat tolerance has been shown to vary depending on the species and even strain of endosymbiont infecting the host. For instance, Gruntenko, Ilinsky, and Adonyeva (2017) demonstrated that the *Wolbachia* strain *w*MelCS increased survival in female *Drosophila melanogaster* after a 4-hour heat shock due to the upregulation of dopamine metabolism, while other strains had a negative impact or no effect. The effects of *Wolbachia* on temperature preferences of *Drosophila* are also strain-dependent, with *w*Ri, *w*Ha, *w*Sh, *w*Tei, *w*MelCS, and *w*MelPop infected flies preferring cooler temperatures and *w*Mau infected flies preferring warmer temperatures than uninfected flies (Arnold *et al*., 2019; Truitt *et al*., 2019; Hague *et al*., 2020). Thus, it is important to understand variation in the capacity for endosymbionts to influence host heat tolerance at both the species and strain levels.

Any impact that an endosymbiont may have on the heat tolerance of its host will also likely depend on its own heat sensitivity. Long-term exposure to heat can reduce or even eliminate endosymbionts in many insects (Corbin *et al*., 2017; Renoz *et al*., 2019). Obligate endosymbionts, which supply critical nutrients to their host, often restrict the thermal tolerance of their host because they are eliminated at temperatures lower than the insect’s upper thermal limit (Wernegreen, 2013; Zhang *et al*., 2019). Facultative endosymbionts, while not essential for host survival, may lose their impact on host heat tolerance if they are also lost at high temperatures. Effects of endosymbionts, including cytoplasmic incompatibility and pathogen protection, can be ameliorated by reductions in endosymbiont density (Hurst *et al*., 2000; Corbin *et al*., 2017; López-Madrigal and Duarte, 2020; Ross *et al*., 2020). As the frequency and duration of heat waves are expected to increase with global warming (WGI IPCC, 2021; He, 2022), there may be unprecedented effects on the persistence of endosymbionts in host populations and on the interactions between endosymbionts and their insect hosts.

Critical and lethal measures of heat tolerance in adults dominate the literature because they are easy to measure and are correlated to species distribution and so can help to predict distributional changes caused by climate change (Kellermann *et al*., 2012; Overgaard *et al*., 2014; Jørgensen *et al*., 2019). However, these traits may overestimate insect heat tolerance as heat sensitivity can vary depending on what life stage is exposed to thermal stress (Kingsolver *et al*., 2011; Lockwood *et al*., 2018; Sales *et al*., 2018) and the effects of heat stress on one life stage may carry over to subsequent life stages (Zhang *et al*., 2015; Porcelli *et al*., 2017; Green *et al*., 2019). For instance, flour beetles exposed to high temperatures (40-42°C) experience greater mortality at pupal and immature-adult life stages than mature adults (Sales *et al*., 2018). Additionally, the impact of temperature on fertility has been overlooked, a trait that can decline at lower than lethal temperatures and is essential for species proliferation (Walsh *et al*., 2019; Parratt *et al*., 2021; Wang and Gunderson, 2022). Parratt *et al*. (2021) and van Heerwaarden and Sgrò (2021) showed that the temperature where *Drosophila* males lost their fertility after experiencing thermal stress during developmental stages or as an adult (upper fertility thermal limit (FTL)) was a better indicator of global *Drosophila* species distributions and laboratory extinction than critical thermal limits. Further, *Wolbachia* are present in immature sperm and can influence the expression of heat-shock proteins (Hsp) in developing larvae (Feder *et al*., 1999; Snook *et al*., 2000). Therefore, understanding how endosymbionts influence heat survival and fertility after heat stress across the life cycle will be crucial for understanding the impact of endosymbionts on insect vulnerability to climate change.

In this paper, we explored the capacity for *Wolbachia* to affect the heat tolerance and fertility of *Drosophila simulans*, a cosmopolitan sister species of *Drosophila melanogaster* (Singh *et al*., 1987). *Wolbachia* is the most common endosymbiont in insects (Weinert *et al*., 2015; Sazama *et al*., 2019), with a wide distribution (Charlesworth *et al*., 2018), and diverse phylogeny (Scholz *et al*., 2020), potentially influencing the thermal tolerance of many insect species. Here, we elucidate the effect of three *Wolbachia* strains that are native to *D. simulans*: *w*Ri, *w*Au, and *w*No. We test the effects of each *Wolbachia* strain on the survival and fertility of *D. simulans* after exposure to heat stress during development or acute heat shock during adulthood. We also address the effect of developmental heat stress on the density of the three *Wolbachia* strains. Understanding the capacity for endosymbionts like *Wolbachia* to influence the upper thermal limit of their host is essential to predicting insect susceptibility to climate change and mitigating the impact of their extinction.

## Materials and Methods

### Sample collection and treatment

*Drosophila simulans* were collected from a range of locations and time points (Table 1). Iso-female lines were established after collection and maintained at a constant 19°C under a 12:12 light:dark cycle on cornmeal-dextrose medium (per litre of water - 73 g cornmeal, 35 g dried yeast, 20 g soy flour, 75 g dextrose, 6 g agar, 16.5 mL nipagin, 14 mL acid mix (546 mL H_2_O, 412 mL propionic acid, 42 mL phosphoric acid)). *Wolbachia* infection and strain identity was confirmed by first amplifying source DNA via PCR using coxA, hcpA, ftsZ, fpbA, and gatB forward and reverse MLST primers (Baldo *et al*., 2006) along with wsp_val primers (Lee *et al*., 2012; Kriesner *et al*., 2013). Source DNA was then sent off for sequencing (Macrogen, Korea).

**TABLE 1.** Summary information of the fly lines used in the paper. *Wolbachia* strain information was sourced from Merçot and Charlat (2004).

| Line | Supergroup | Mitochondrial<br>haplotype<br>association | CI<br>phenotype | Source location | Source<br>date |
| --- | --- | --- | --- | --- | --- |
| wRi | A | sII | *mod + resc*<br>+ | Yarra Valley,<br>VIC, Australia | 2019 |
| wAu | A | sII | *mod -* | Perth Hills, WA,<br>Australia | 2018 |
| wNo | B | sI | *mod + resc*<br>+ | Noumea, New<br>Caledonia | Stock (est.<br>1980s) |
| Naturally<br>uninfected | - | - | - | Yarra Valley,<br>VIC, Australia | 2019 |

*Wolbachia*-uninfected (cured) lines were created by rearing *Wolbachia*-infected lines on cornmeal-dextrose medium containing tetracycline antibiotics (0.03%) for two generations (Richardson *et al*., 2016). Flies were reared in the absence of antibiotics for two additional generations prior to backcrossing. Removal of *Wolbachia* was verified using quantitative PCR (qPCR) by testing for the presence of the *Wolbachia* surface protein (*wsp*) gene (see *Wolbachia density* section below for qPCR protocol details). For each line, 16 adult flies were randomly selected for testing (irrespective of sex) and two technical replicates were generated.

Cured and *Wolbachia*-infected lines were then backcrossed with a naturally uninfected line sourced from the same location and time as the *w*Ri line (Table 1). This was done to control for differences in nuclear genetic background, though differences in mitochondria were retained due to maternal transmission. Twenty unmated females from each cured and *Wolbachia* infected line were mated with twenty males from the naturally uninfected line for four generations. Flies were sexed using CO_2_ anaesthesia under a dissecting microscope and rested for 48 hours before crossing. *Wolbachia* infection status was confirmed prior to starting the experiments with qPCR. The six backcrossed lines used in the experiments are henceforth referred to as *w*Au, *w*Ri, *w*No (*Wolbachia* infected), *w*Au.tet, *w*Ri.tet, and *w*No.tet (tetracycline-cured).

### Heat tolerance assays

#### Treatment of experimental lines

Flies were maintained at constant 25°C, corresponding to the midpoint of the optimum developmental thermal range for *D. simulans* (David *et al*., 2005; Austin and Moehring, 2013), under a 12:12 light:dark cycle for more than four generations before assessing heat tolerance. Prior to any experiment, fly density was partially controlled via short lays (3-4 hours) in the parental generation to avoid stress from overcrowding. Sexes were separated under CO_2_ then allowed to recover on fresh media for at least 48 h prior to any experiment (MacMillan *et al*., 2017).

#### Heat stress assays

Heat tolerance in infected and uninfected lines was assessed by estimating both survival and fertility after heat stress during development, or after an acute heat shock during the adult stage.

#### Upper egg-to-adult developmental lethal thermal limit

The upper developmental lethal thermal limit (devLTL) is the uppermost temperature at which flies successfully develop from egg to adult (Petavy *et al*., 2001; Overgaard *et al*., 2014; van Heerwaarden and Sgrò, 2021). Developmental LTLs were assessed by rearing eggs in PHCbi controlled temperature (CT) cabinets set to different fluctuating temperature regimes (28, 29, 30, 31, or 32 ±3°C) with a 12:12 light:dark cycle (see van Heerwaarden and Sgrò (2021) for regime). The temperature range was chosen based on data from van Heerwaarden and Sgrò (2021) and refined by a pilot experiment (Supplementary Figure S1).

In the parental generation, six to seven-day old adult flies were placed in inverted 500 mL containers (lay cages) that had a thin layer of fly food coloured with food dye in the lid. Live yeast was sprinkled onto the media to stimulate oviposition and the flies were allowed to lay overnight at constant 25°C. Twenty eggs were picked from the media under a dissecting microscope and placed in a vial containing 20 mL of cornmeal media. Twenty vials were set up for each line and temperature and placed in the CT cabinets to allow the flies to develop. Once the flies had started to emerge, they were left for five days to ensure all viable adults had eclosed. The vials were then placed at 4°C to knock the flies out before scoring viability. Flies were only scored as viable if they had completely emerged from their pupal case. The experiment was blocked across two generations, with 10 vials of 20 eggs scored per generation for each line.

#### Male upper developmental fertility thermal limit

The upper developmental fertility thermal limit (devFTL) is the uppermost temperature that a developing fly can be exposed to and still produce offspring when fully matured (van Heerwaarden and Sgrò, 2021). To assess the devFTL of adult male flies developed under warming conditions, eggs were reared in CT cabinets with fluctuating temperature regimes (28, 28.5, 29, 29.5, or 30°C ±3°C) with a 12:12 light:dark cycle (van Heerwaarden and Sgrò, 2021).

Adult flies were set up in lay cages as described previously. Forty to fifty eggs were then cut from the fly media and transferred to vials containing 20 mL of cornmeal media before placing them in the CT cabinets to allow the flies to develop. After emerging, flies were sexed and males were placed on fresh food and returned to the CT cabinets. After five days, 20 males from each line and temperature were placed in individual vials with two unmated females from the same line. Unmated females had been sexed five days prior and maintained at constant 25°C. Vials were placed back in the relevant CT cabinets in which the males had been reared and flies were allowed to mate and lay eggs for seven days. Males were scored as sterile if larval activity could not be detected in the vial under a dissecting microscope after seven days of mating.

#### Wolbachia density in adult male flies

Adult male flies were collected 24-48 hours after eclosing following exposure to the temperatures in the viability experiment above, except for 32°C which had zero viability. Flies reared from egg to adult at 25°C as a control were also collected. *w*No flies reared at 19°C were also included to validate the lower density of this strain, though these flies were 10 days old and collected 2 months after the other flies. Each treatment included 16 biological replicates, each with 2-3 technical replicates. Genomic DNA was extracted from whole adult male bodies using a Chelex-based method in 1.7 mL tubes (Richardson *et al*., 2016). Tubes contained 150 µL of 5% Chelex solution and 2.5 µL proteinase K and were incubated for 1 h at 65°C followed by 10 min at 90°C before being diluted in 96-well plates (10 µL supernatant to 90 µL purified water in each well). The *w*Ri and *w*Au lines were blocked across each qPCR run to account for run differences. *w*No infected flies were analysed separately due to the low density of *Wolbachia* (Osborne *et al*., 2012). *Wolbachia* density was estimated by assessing the ratio of the single copy *wsp* gene (*Wolbachia*) to the *RpL40* reference gene (*Drosophila*) using quantitative PCR (qPCR) via the Roche LightCycler480 system, according to Lee *et al*., (2012) (see Supplementary Table S1 for primer sequences).

#### Male heat shock upper lethal thermal limit

The male adult heat shock upper lethal thermal limit (hsLTL) represents the highest temperature before irreversible and fatal damage occurs in adults after a heat shock (Cowles and Bogert, 1944; Terblanche *et al*., 2011; Parratt *et al*., 2021). To assess the male adult hsLTL, 0.5 mL microcentrifuge tubes containing five adult male flies (five days old) were placed in Biometra TRIO thermal cyclers with base temperatures set to constant 36.5, 37, 37.5, 38, 38.5, or 39°C for a 1-hour heat shock (Kong *et al*., 2016). Ten tubes were set up for each line and temperature (50 flies per line per temperature). The thermal cycler lid temperature only provided 1.0°C resolution, so was set to the nearest whole integer of each base temperature rounded down (36, 37, 37, 38, 38, and 39°C respectively). The temperature range was initially estimated from Parratt *et al*. (2021) and refined via a pilot experiment to suit the shorter heat shock duration (Supplementary Figure S2). After 1 hour of heat shock, the microcentrifuge tubes were placed in vials containing 20 mL of cornmeal media and the flies were allowed to recover for 24 hours at constant 25°C. Each fly was scored as surviving if any movement could be detected. The experiment was blocked across two runs in the same day, with each run containing five microcentrifuge tubes of flies from each line.

#### Male adult heat shock upper fertility thermal limit

The male adult heat shock upper fertility thermal limit (hsFTL) was assessed on surviving male flies 48 hours after heat shock from the hsLTL experiment above. Twenty adult males (10 flies randomly selected from each experimental block) per line, per temperature, were placed in individual vials containing 20 mL of cornmeal media. Each male was paired with two adult unmated females from the same line that had been sexed 6-7 days prior and maintained at constant 25°C. The mating status of these females was confirmed by checking larval activity in the holding vials prior to mating. Vials were placed at 25°C and flies were allowed to mate and lay eggs for seven days. Male sterility was scored as described previously.

### Statistical analysis

#### Wolbachia density in adult male flies

*Wolbachia* density was estimated using the formula:

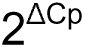

Where delta Cp is the difference between the *wsp* and *RpL40* primer Cp values, averaged across two to three consistent replicates. Results were analysed using a polynomial linear regression model (*stats* package, R Core Team, 2022) and a two-way ANOVA (*car* package, Fox and Weisberg, 2019) with the response variable as *Wolbachia* density and explanatory variable as temperature. Model fit was assessed by comparing the adjusted R^2^ value of different models and model assumptions were verified by plotting the regression and assessing the diagnostic information.

#### Upper developmental lethal thermal limit, male developmental fertility thermal limit, and heat shock lethal thermal limit

Thermal limits were calculated using a dose response model (drm) that was created using the *drc* package (Ritz and Strebig, 2016). The upper heat shock lethal thermal limit and the developmental fertility thermal limit were determined by the temperature at which 50% of flies were still alive (hsLTL_50_) or still fertile (devFTL_50_), while the upper developmental lethal thermal limit was determined by the temperature at which the maximum possible viability was halved (devLTL_50_) – this is because the maximum viability never reached 100%. Model fit was assessed using the *mselect* function (*drc* package) with the AIC selection method. The model response variable was the proportion of viable/fertile/surviving adults, and the explanatory variables were temperature and infection status. devLTL_50_/devFTL_50_/hsLTL_50_ values were generated using the *ED* function (*drc* package) with interval set to delta and compared using the *EDcomp* function (*drc* package). A two-way ANOVA (*car* package) of a generalised linear model (*stats* package) was implemented to assess whether block effects influenced each trait. The response variable was the proportion of viable flies/ fertile males/ surviving males and the explanatory variables were block number, temperature, and infection status. Where block effects were significant, we included Tukey’s post-hoc tests (*stats* package) and additional figures in the supplementary section to display the results by block.

#### Male adult heat shock upper fertility thermal limit

Statistical analysis was performed using a two-way ANOVA (*car* package) on a generalised linear model with family set to binomial and weight set to the total number of flies measured. The response variable was the proportion of fertile males, and the explanatory variables were block number, temperature, and infection status and their interactions. hsFTL_50_ values could not be generated using the same method outlined above as male fertility after heat shock did not follow a standard dosage response curve.

#### Figure creation

Graphs were generated using the *ggplot2* package (Wickham, 2016) and arranged using the *ggarange* function (*ggpubr* package, Kassambara, 2023).

## Results

### The developmental lethal thermal limit is reduced by *Wolbachia* infection

We first examined whether each *Wolbachia* infection affected the 50% upper developmental lethal thermal limit (devLTL_50_) of flies reared at fluctuating developmental temperatures ranging from 28-32 ± 3°C. Infection with *w*Au or *w*Ri significantly reduced the devLTL_50_ compared to the respective cured lines (Estimated ratio of lethal limit: *w*Au: Δ = -0.33°C, t = -3.99, p = <0.001, *w*Ri: Δ = -0.18°C, t = -2.32, p = 0.021) (Figure 1). *w*No infection also reduced the devLTL_50_, but the difference was not significant (Estimated ratio of lethal limit: Δ = 0.17°C, t = -1.91, p = 0.056) (Figure 1c). We found a significant block effect in the *w*Au line (ANOVA: *χ*^2^_(3)_ = 7.87, p = 0.049) that was driven by differences between blocks 3 vs 1 and 3 vs 2, though the direction of the effect of *Wolbachia* in each block was consistent (Supplementary Table S3, Supplementary Figure S3).

**FIGURE 1.**
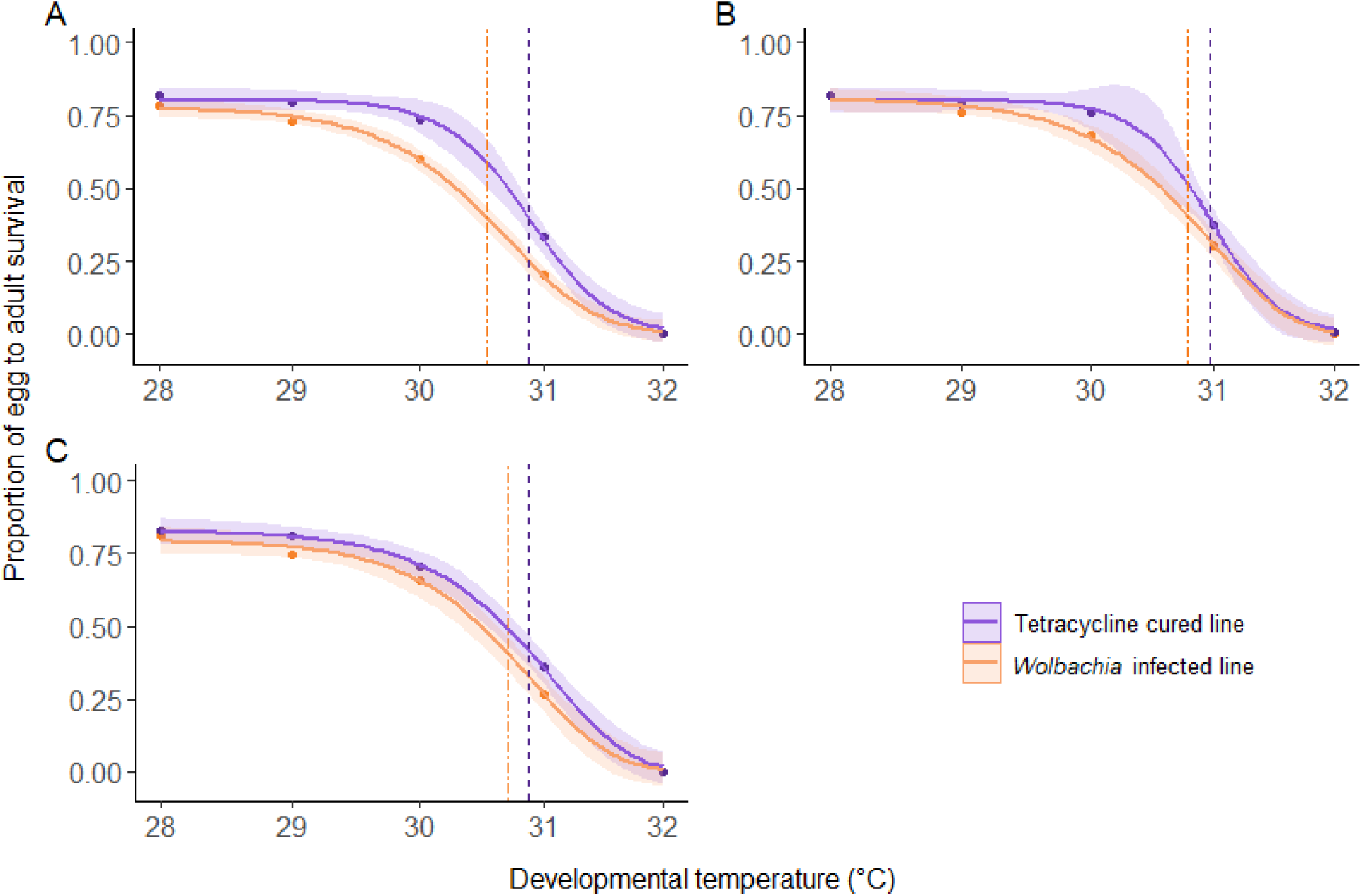
The proportion of surviving *D. simulans* reared from egg to adult at different fluctuating developmental temperatures for the (a) *w*Au, (b) *w*Ri, and (c) *w*No lines. Points show the mean viability at each temperature and lines were generated using predictions from dosage response models with shading representing the 95% confidence interval of the predicted means. Purple lines represent tetracycline cured lines and orange lines represent *Wolbachia*-infected lines. Dashed vertical lines represent the devLTL_50_ values for the fly lines in their respective colours.

### The fertility thermal limit of adult males after developmental heat stress is reduced by *w*No *Wolbachia* infection

We next assessed the impact of *Wolbachia* infection on the 50% upper developmental fertility thermal limit (devFTL_50_) of adult males developed under fluctuating temperatures between 28-30 ±3°C. *w*Ri infection did not significantly change the devFTL_50_ compared to *w*Ri.tet (Estimated ratio of fertility limit: Δ = 0.05°C, t = -0.31, adj. p = 0.760) (Figure 2). However, infection with *w*No and *w*Au reduced the FTL_50_, though the latter was not significant (Estimated ratio of fertility limit: *w*No: Δ = -0.17°C, t = 1.87, p = 0.015; *w*Au: Δ = -0.14°C, t = 1.87, adj. p = 0.063).

**FIGURE 2.**
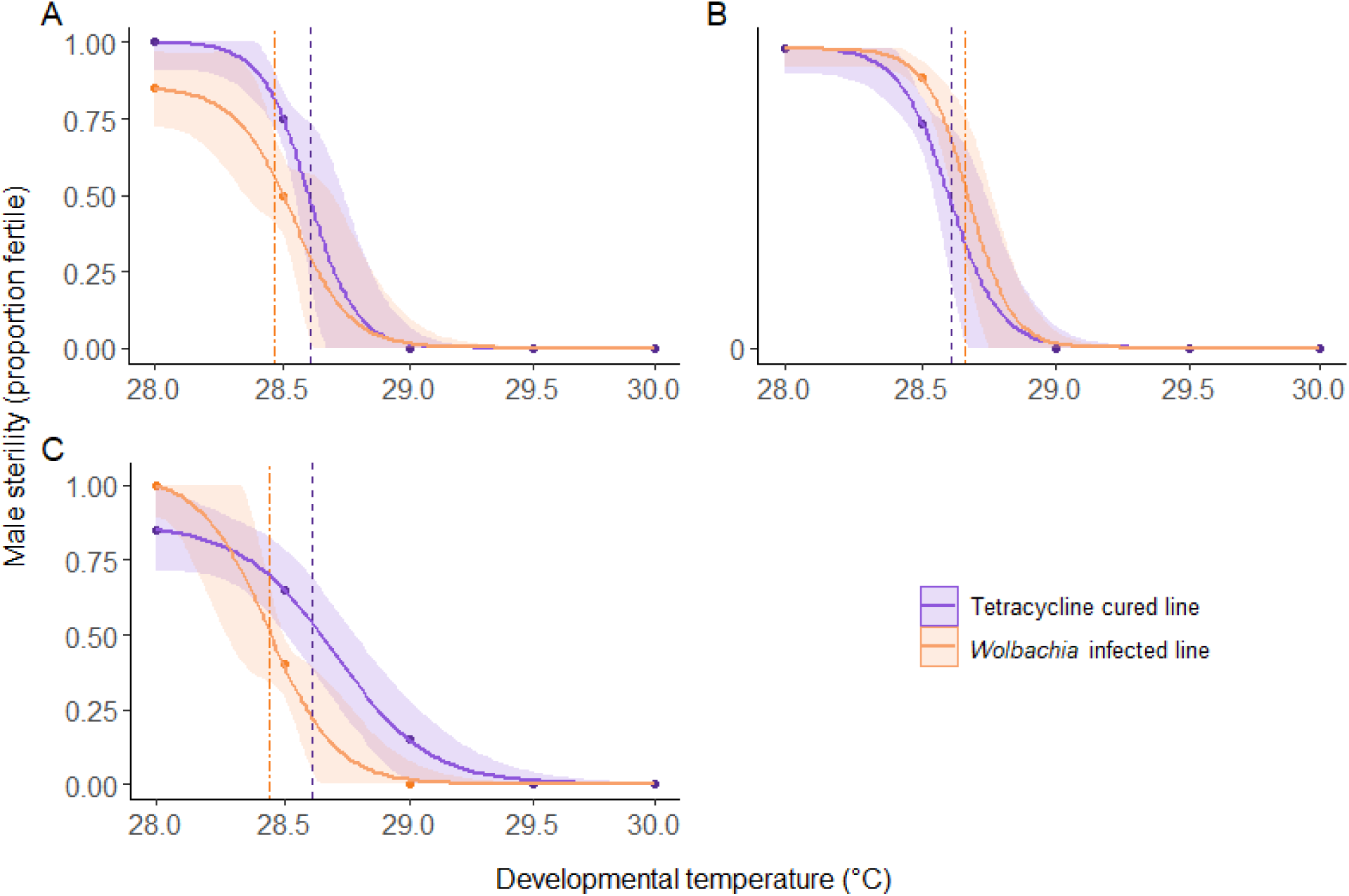
Male sterility (measured as the proportion of fertile flies) in *D. simulans* reared from egg to adult at different fluctuating temperatures for the (a) *w*Au, (b) *w*Ri, and (c) *w*No lines. Points show the mean fertility at each temperature and lines were generated using predictions from dosage response models. Purple lines represent tetracycline cured lines and orange lines represent *Wolbachia* infected lines. Dashed vertical lines represent the devFTL_50_ values for the fly lines in their respective colours.

### *Wolbachia* density is reduced by temperatures below the viability thermal limit

We wanted to understand the changes in density of the three *Wolbachia* strains in flies that had developed from egg to adult under different fluctuating temperatures (28-31 ± 3°C and a control at constant 25°C). Developmental temperature had a significant effect on *Wolbachia* density for each strain (ANOVA: *w*Au: F_(1, 75)_ = 8.36, p = 0.005; *w*Ri: F_(1, 76)_ = 4.06, p = 0.05; *w*No: F_(1, 86)_ = 76.23, p = <0.001) (Figure 3). *Wolbachia* density was highest between 25-29°C for *w*Au while density peaked between 28-29°C for *w*Ri (Figures 3a and 3b, Supplementary Table S5). For *w*No, density reduced considerably between 19 and 25°C (Tukey’s HSD: p = <0.001) but there were no pair-wise differences between each successive temperature above 25°C (Figure 3c, Supplementary Table S5).

**FIGURE 3.**
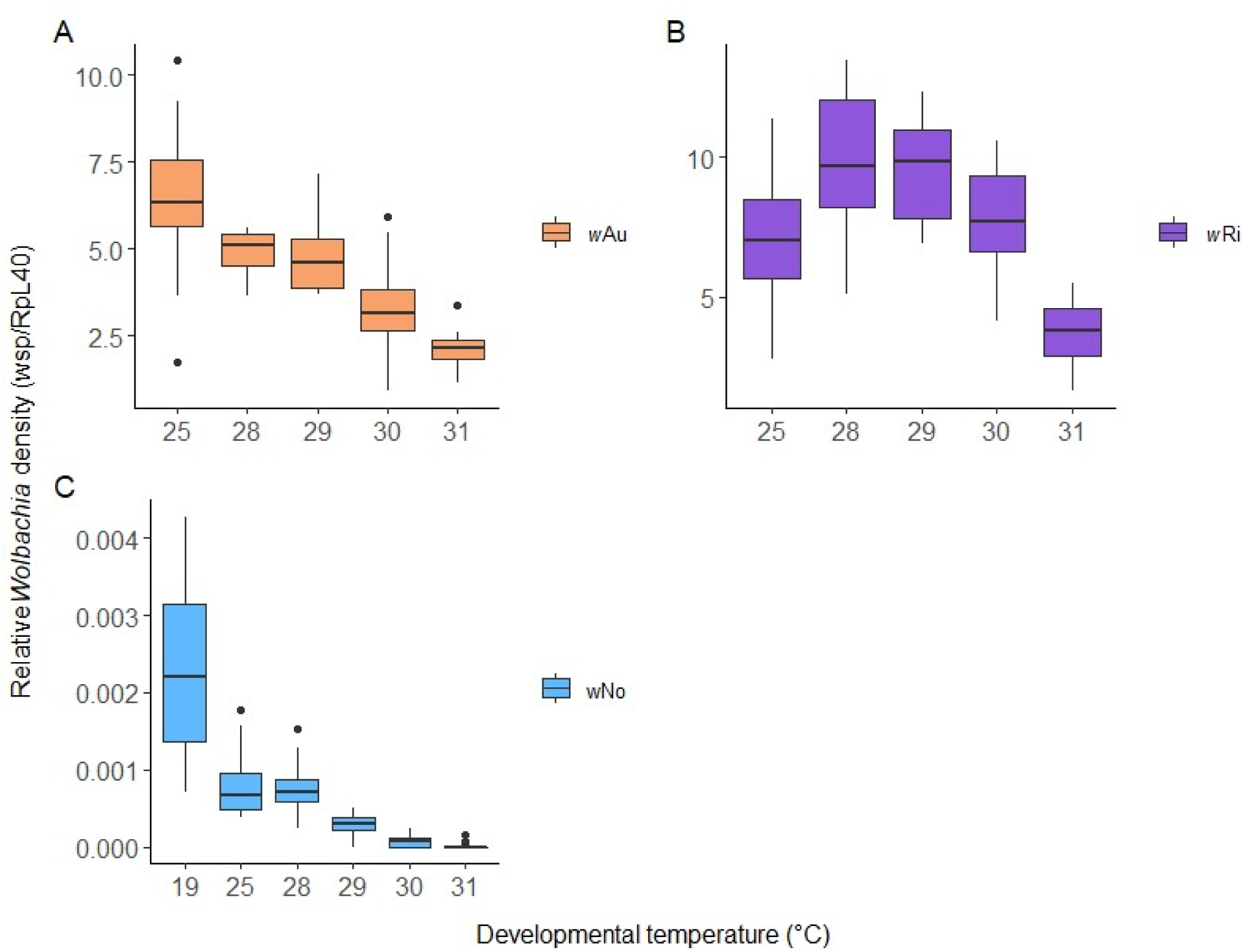
*Wolbachia* density for (a) *w*Au, (b) *w*Ri, and (c) *w*No strains infecting male *D. simulans* reared from egg to adult at different fluctuating developmental temperatures. Horizontal black bars represent means, whiskers show data range, and points show outliers. For panel C, flies from 19°C were reared under different experimental conditions (see Materials and Methods section).

### The adult heat shock lethal thermal limit is reduced in male flies infected with *w*No

We wanted to determine the effect of *Wolbachia* infection on the 50% upper lethal thermal limit (hsLTL_50_) of male flies exposed to an acute heat shock at a range of static temperatures between 36.5-39°C. Flies infected with *w*No had a significantly lower hsLTL_50_ than the *w*No.tet line (Estimated ratio of lethal limit: Δ = -0.16°C, t = -4.26, p = <0.001) (Figure 4). *w*Au-infected flies also showed a lower hsLTL_50_, though the effect was not significant (Estimated ratio of lethal limit: Δ = -0.06°C, t = -1.76, p = 0.079), but *w*Ri-infection did not impact the hsLTL_50_ (Estimated ratio of lethal limit: t = -0.50, p = 0.616). There was a significant block effect in the *w*No line (ANOVA: *χ*^2^_(1)_ = 4.10, p = 0.043), with block one having a lower average survival than block 2, though the direction of the effect of *Wolbachia* in each block was consistent (Supplementary Figure S4).

**FIGURE 4.**
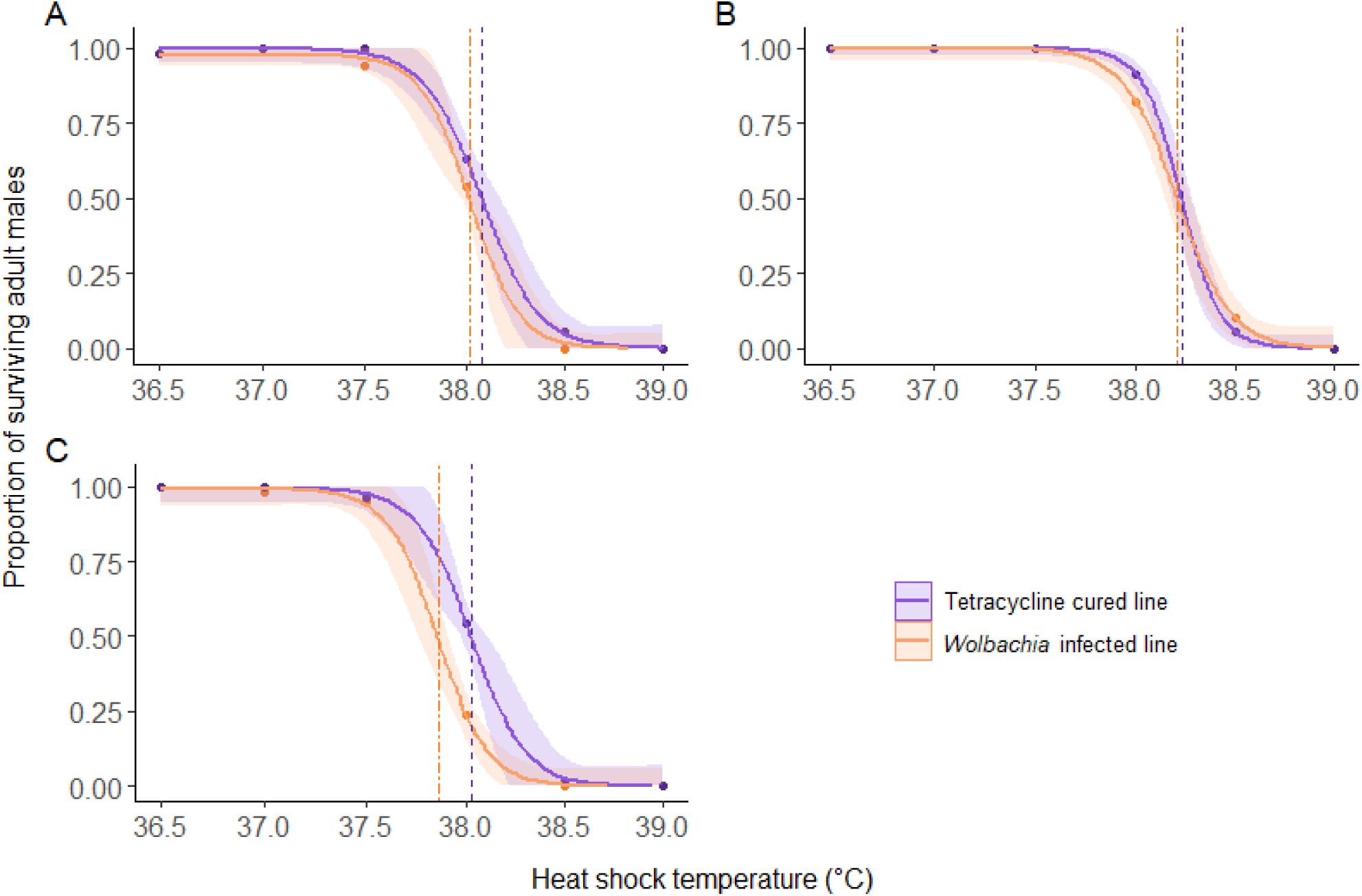
The proportion of surviving adult male *D. simulans* exposed to a static 1-hour heat shock for the (a) *w*Au, (b) *w*Ri, and (c) *w*No lines. Points show the mean survival at each temperature and lines were generated using predictions from dosage response models with shading representing the 95% confidence interval of the predicted means. Purple lines represent tetracycline cured lines and orange lines represent *Wolbachia*-infected lines. Dashed vertical lines represent the hsLTL_50_ values for the fly lines in their respective colours.

### The adult male heat shock fertility thermal limit is reduced in flies infected with *w*No

After exposing male flies to acute static heat shock between 36.5-38°C, we assessed whether *Wolbachia* infection impacted male sterility over a seven-day period. Neither *w*Au nor *w*Ri *Wolbachia* infections impacted the proportion of fertile males after heat shock (ANOVA: *w*Au: *χ*^2^_(1)_ = 1.23 p = 0.267, *w*Ri: *χ*^2^_(1)_ = 0.55, p = 0.453) (Figure 5). However, *w*No infection significantly reduced the proportion of fertile males (*w*No: *χ*^2^_(1)_ = 5.66, p = 0.018) (Figure 5). Increased temperature generally increased sterility (ANOVA: X^2^_(1)_ = 18.66 p = <0.001), which was the case for *w*Ri and *w*Au fly lines (ANOVA: *w*Au: *χ*^2^_(1)_ = 13.91 p = <0.001, *w*Ri: *χ*^2^_(1)_ = 5.62, p = 0.017). However, there was no significant impact of temperature on the fertility of the *w*No line (ANOVA: *w*No: *χ*^2^_(1)_ = 1.84, p = 0.170). Fertility remained high across all fly lines up to the survival thermal limit of 39°C, though high mortality at 38.5°C prevented comparisons of fertility at this temperature. Notably, the *w*Au.tet line saw a significant drop in fertility between 37.5-38°C (Tukey’s HSD: Δ = -0.40%, p = 0.003), but this was not significantly different from the *w*Au line at 38°C (Tukey’s HSD: p = 0.112).

**FIGURE 5.**
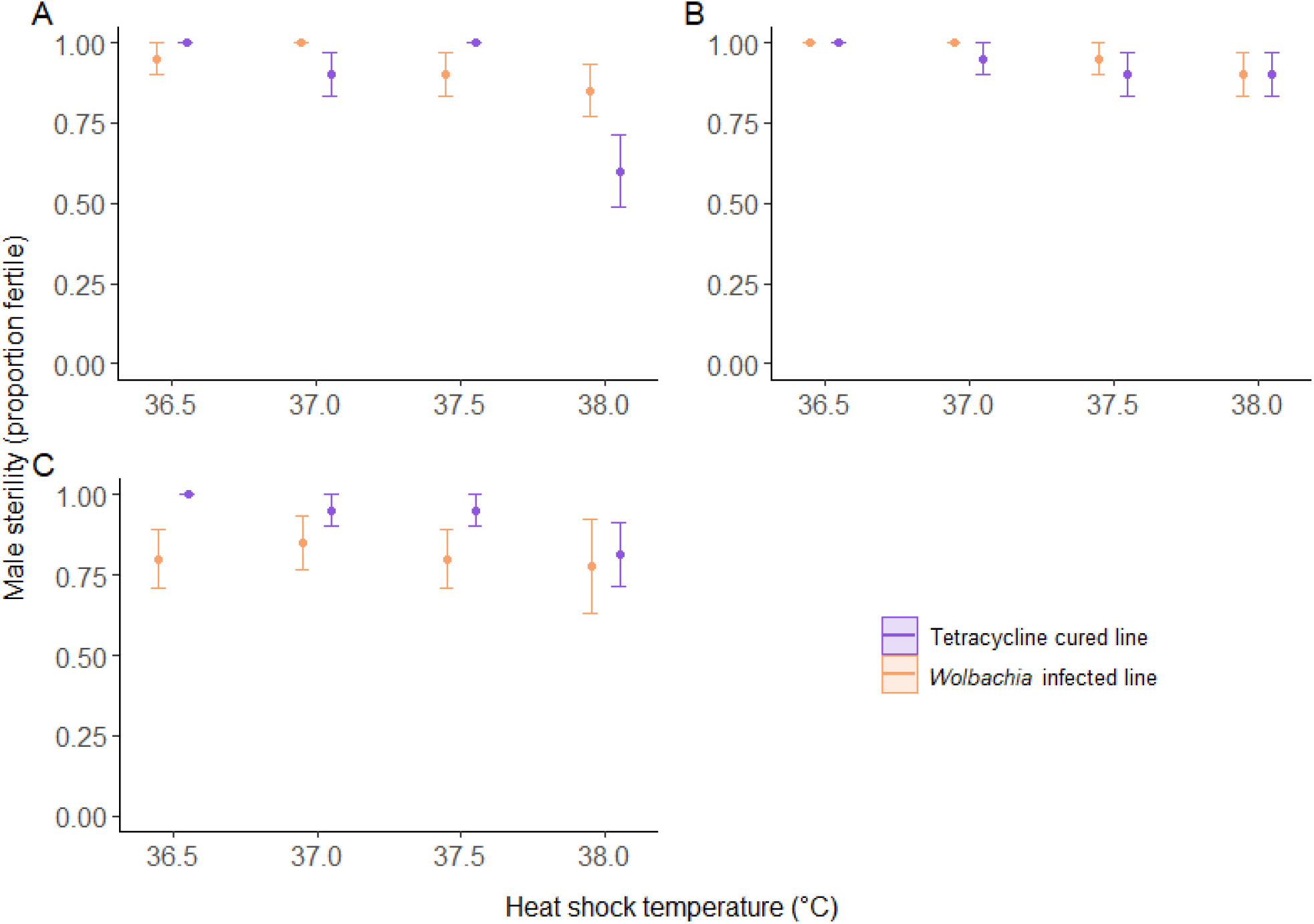
The proportion of fertile adult male *D. simulans* exposed to a static 1-hour heat shock for (a) *w*Au, (b) *w*Ri, and (c) *w*No lines. Points represent the mean fertility at each temperature and bars show the standard error of the mean.

## Discussion

Current trait-based estimates of climate change vulnerability ignore the impact of endosymbionts on fitness, despite evidence demonstrating the capacity for endosymbionts to influence the heat tolerance of their host (Wernegreen, 2013; Corbin *et al*., 2017; Dunn, 2017; Hector *et al*., 2022). Further, measures of upper LTLs at the adult life stage ignore the impact of heat on earlier stages and the fertility of surviving adults. Fertility can decline at much lower temperatures than the upper LTL and is emerging as an important trait in assessing insect species distribution and climate change vulnerability (Walsh *et al*., 2019; Parratt *et al*., 2021; van Heerwaarden and Sgrò, 2021). Here, we demonstrate that *Wolbachia* infection has a small, negative effect on the survival of *D. simulans* when exposed to heat stress during development or as an adult, though the size of the effect is dependent on the *Wolbachia* strain. Further, the fertility of adult males exposed to heat stress at both juvenile and adult life stages is also reduced in a strain-specific manner.

The strain and trait-specific way by which *Wolbachia* influences thermal tolerance highlights the potential for misleading conclusions when only one trait is considered in assessing heat tolerance. Here, we found that the egg-to-adult lethal thermal limit (devLTL) of the flies was reduced by both *w*Au and *w*Ri *Wolbachia* infections, though not by *w*No infection. In contrast, the developmental fertility thermal limit (devFTL) was only reduced by *w*No infection and not by *w*Au or *w*Ri infections. Without considering both survival and fertility, which are essential for fitness, we would not have identified that all three *Wolbachia* strains have a small, negative effect on heat tolerance under developmental heat stress. The results support the suggestion by some authors of moving towards a multi-trait view of heat tolerance (Blackburn *et al*., 2014; Walsh *et al*., 2019). There is also the possibility that small effects in one trait may be offset by effects in another trait, or that the effects compound to be quite substantial, which would only be discoverable by assessing multiple traits. We found that adult males became sterile before they approached the devLTL, meaning the reduced egg-to-adult survival caused by the *Wolbachia* infection may not be ecologically important because the surviving adult male flies would not be able to reproduce anyway. However, we did not explore the capacity for fertility to recover after exposure to developmental heat stress, as flies were mated at the same stressful developmental temperatures, meaning survival of the juvenile offspring would still be relevant. Furthermore, like many insect species, *D. simulans* has overlapping generations in the wild, so reductions in developmental survival may still be important, especially when there is heterogeneity across microhabitats. Nonetheless, the results demonstrate the importance of assessing fertility after survival when insects are exposed to heat stress during development and of considering *Wolbachia* strain differences to better predict insect vulnerability to climate change.

The varying effects of the *Wolbachia* strains under developmental heat stress may be due to phylogenetic differences; one impact of which is differences in tissue distribution and density (Merçot and Charlat, 2004; Osborne *et al*., 2012). *w*No, which belongs to supergroup B, generally maintains a very low whole-adult density and is undetectable in some tissues. In contrast, *w*Au and *w*Ri, which belong to supergroup A, have a high abundance (Merçot and Charlat, 2004). Here, we observed that *w*Au and *w*Ri maintained high densities up to the devLTL while *w*No reduced in density by an order of magnitude at the devFTL and was effectively eliminated at the devLTL. While density has been linked to phenotypes affected by *Wolbachia*, including antiviral protection (Osborne *et al*., 2012; Martinez *et al*., 2014), there is evidence that some strains with low densities can still produce strong phenotypic effects (Richardson *et al*., 2019). The lower temperatures that maintained the low-density *w*No infection at the devFTL may have facilitated the negative effect of *Wolbachia* on adult fertility, while the higher temperatures at the devLTL may have attenuated the effect of *w*No infection on egg-to-adult survival. Osborne *et al*. (2012) showed that *w*No density is highest in the testes of *Drosophila* compared to other tissues and is comparable to the density of *w*Ri (though *w*Au density is higher). While we did not test the effects of temperature on the density of *Wolbachia* in different tissues, the higher relative density of *w*No in the testes compared to other strains may help to explain these strain and trait-specific effects. We acknowledge that a reduction in *Wolbachia* density beyond the devFTL may not be ecologically relevant. However, any decline in *Wolbachia* density below the upper fertility thermal limit may reduce impacts of *Wolbachia* on heat tolerance, particularly if this leads to decreased maternal transmission.

The male adult heat shock lethal thermal limit (hsLTL) and fertility thermal limit (hsFTL) were reduced by *w*No infection, but not by *w*Au or *w*Ri infections. The broader literature also shows diverse effects of strain on adult survival following heat shocks (Brumin *et al*., 2011; Heyworth and Ferrari, 2015, 2016; Gruntenko *et al*., 2017; Zhu *et al*., 2021). For all fly lines, males were not sterilised by heat shock temperatures below the hsLTL, though there was a general increase in sterility with higher temperatures, and this was not impacted by *Wolbachia* infection, which has not previously been shown. This result is consistent with data for *D. simulans* from Parratt *et al*. (2021) and confirms that there is little sublethal effect of heat shock on male sterility prior to death for this species. Nonetheless, it is possible that the fertility of *Wolbachia*-infected and cured flies recovered at different rates after heat shock in the seven days that the male flies mated with females at a benign temperature (Jørgensen *et al*., 2006; Walsh *et al*., 2019), or there was a delayed impact of heat stress on fertility (Parratt *et al*., 2021). Although we chose to focus on the effect of *Wolbachia* on male fertility after heat stress, future studies could also explore whether *Wolbachia* infection influences the survival of sperm stored in the spermatheca of mated females (Sales *et al*., 2018; Walsh *et al*., 2022).

Here, we demonstrated that infection with *Wolbachia* can have a negative impact on the survival and fertility of *D. simulans* when exposed to heat stress during development or heat shock as an adult. While the effects were small and, in the case of developmental survival, may be eclipsed by the sterility achieved at lower temperatures, the effect of *w*No infection on both survival and fertility may compound to have a substantial effect on fitness. The diversity of *Wolbachia* strains and the hosts they inhabit, as well as the variability with which *Wolbachia* influences different heat tolerance traits, highlight the need to consider the presence of endosymbionts when measuring heat tolerance in future studies.

## Supplementary Materials

**Figure S1.**
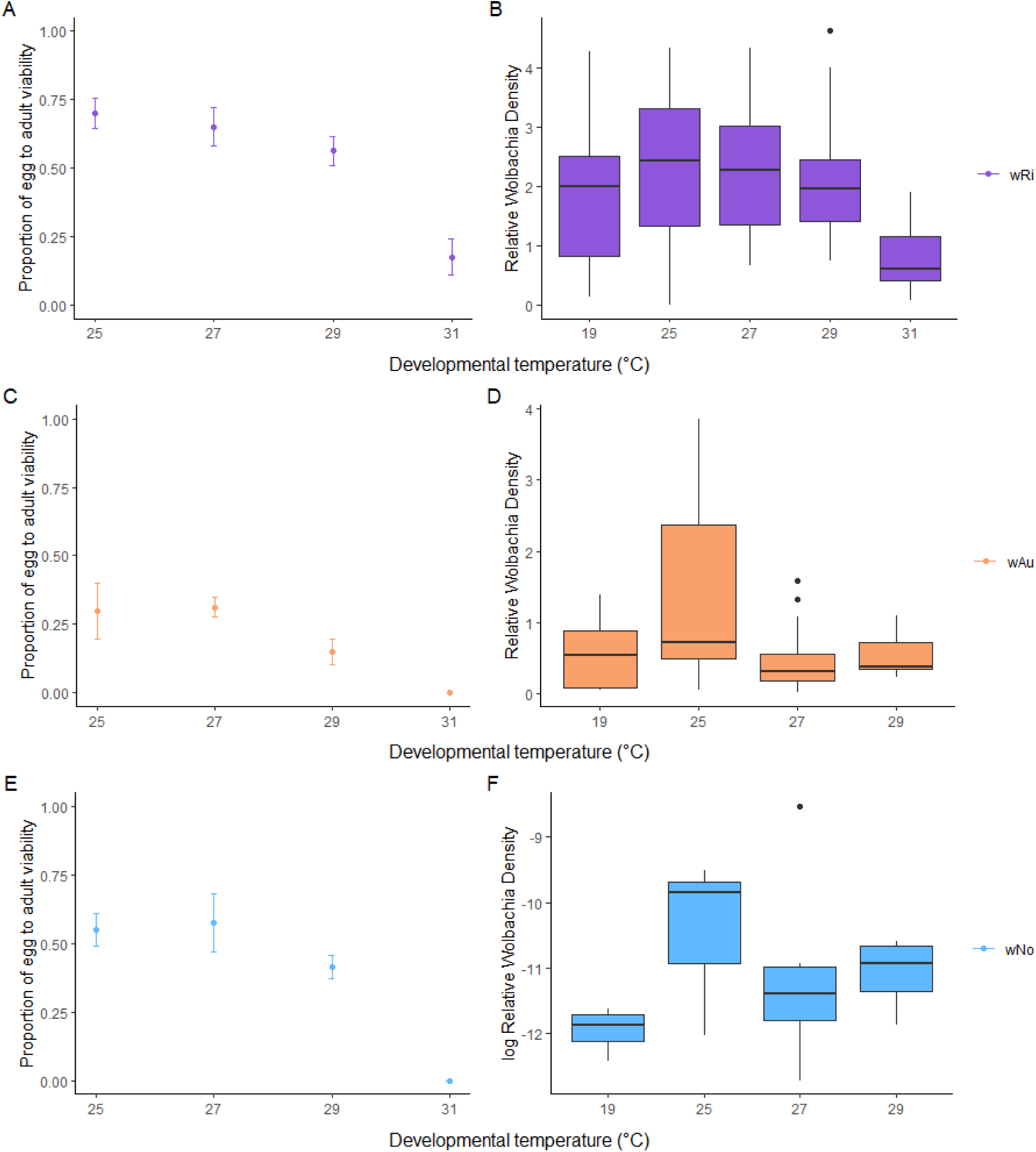
Pilot experiments on *w*Ri- (a-b), *w*Au- (c-d), or *w*No- (e-f) *Wolbachia* infected *D. simulans*. Left: the proportion of surviving flies reared from egg to adult at different fluctuating developmental temperatures. Right: *Wolbachia* density in surviving adult flies after developmental heat treatment.

**Table S1.**
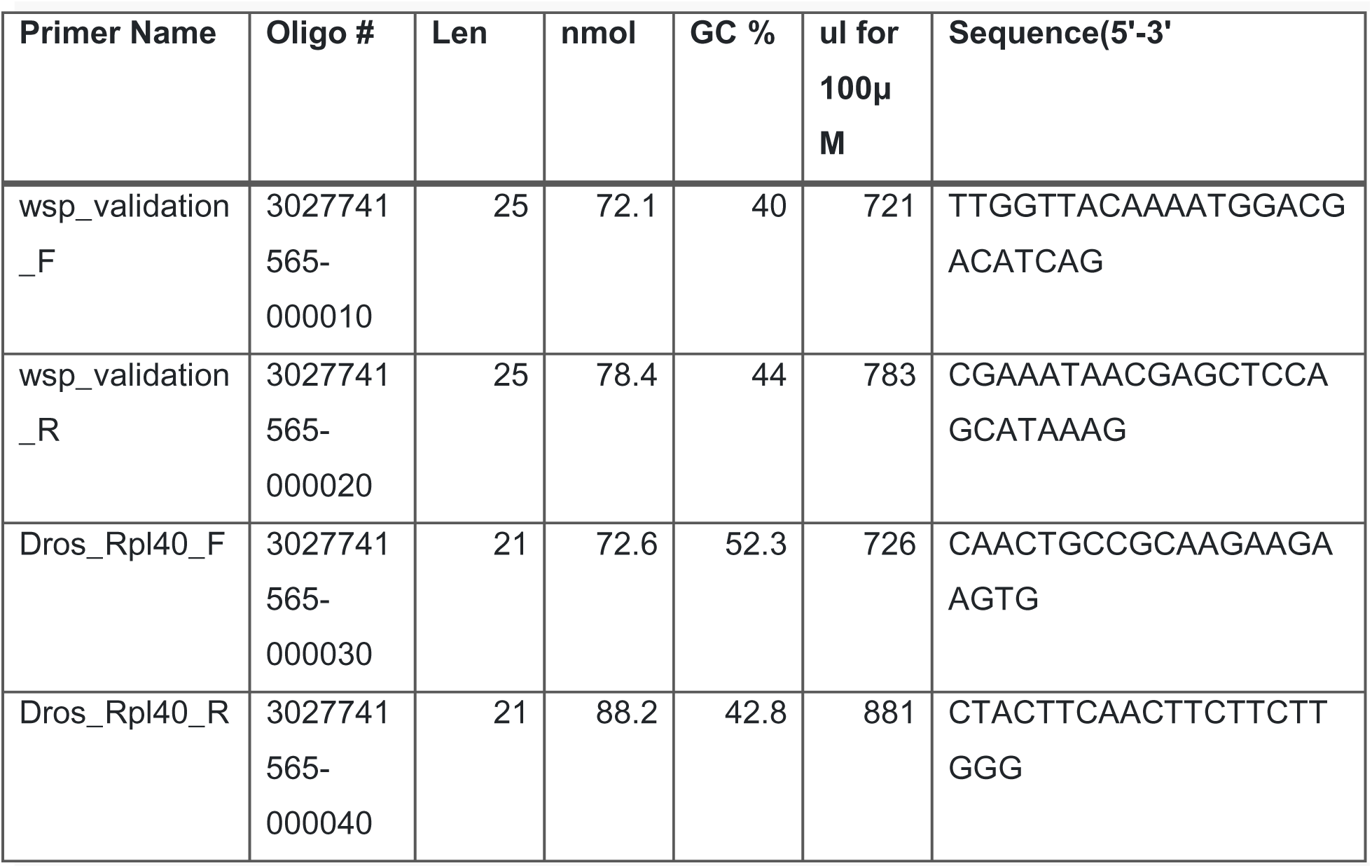
Validation primer information for quantitative PCR work. Primers sourced from Sigma-Aldrich in August 2021.

**Figure S2.**
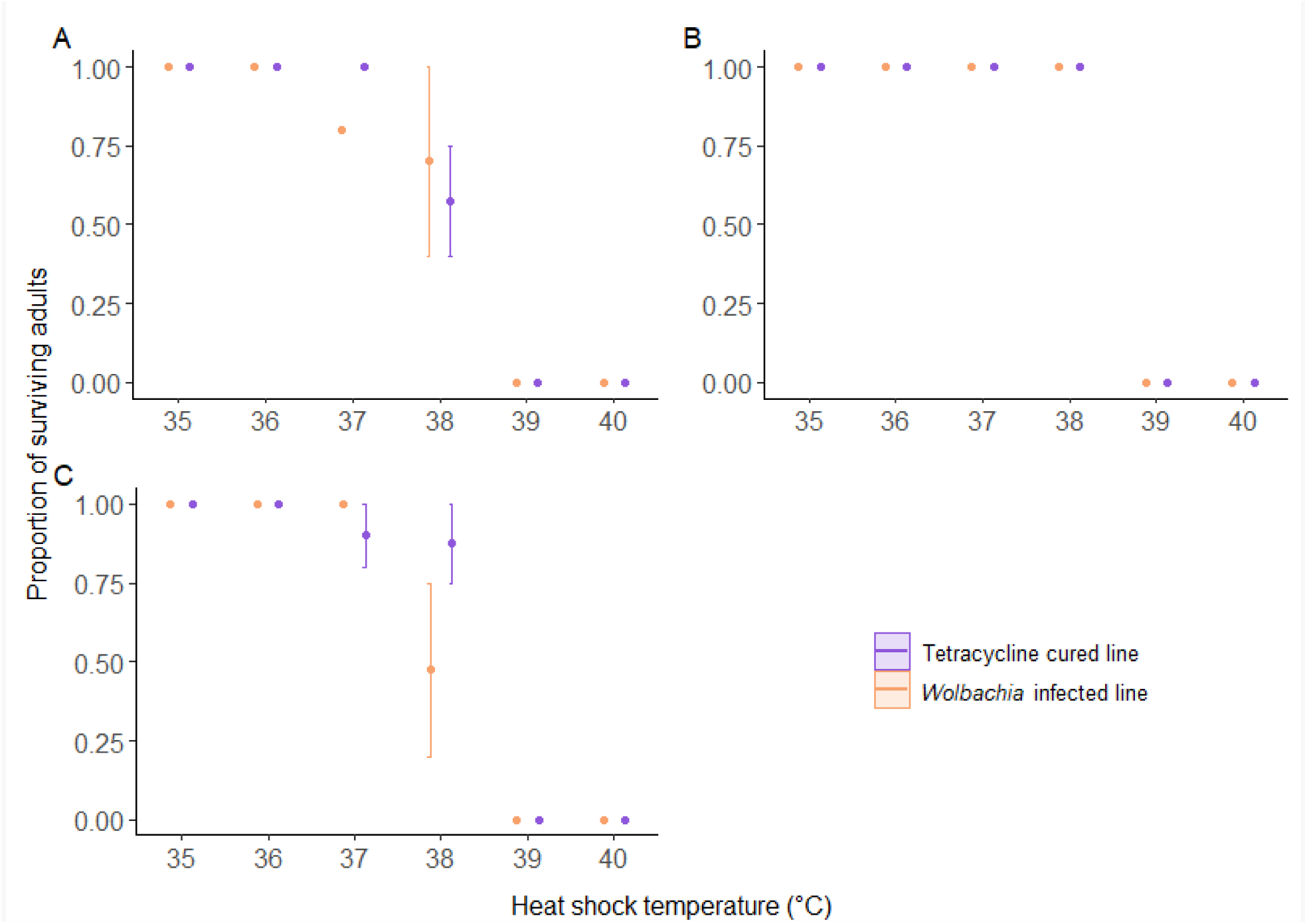
Pilot experiment showing the proportion of surviving adults exposed to a 1 h heat shock. Flies were infected with (a) *w*Au, (b) *w*Ri, or (c) *w*No *Wolbachia*.

**Table S2.**
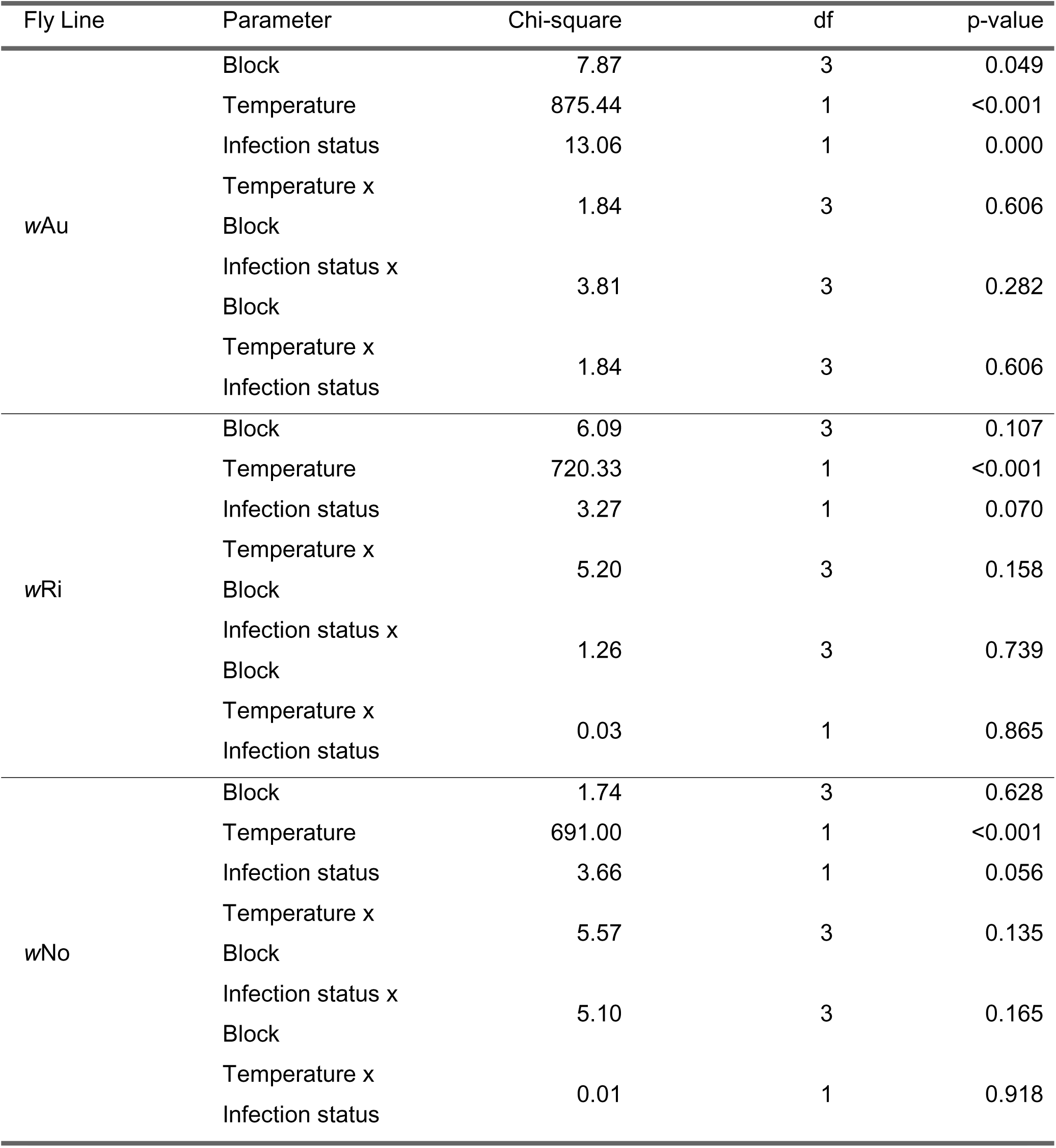
Two-way ANOVA assessing the impact of experimental block, fluctuating developmental temperature, *Wolbachia* infection status, and their interactions on developmental egg-to-adult survival.

**Table S3.**
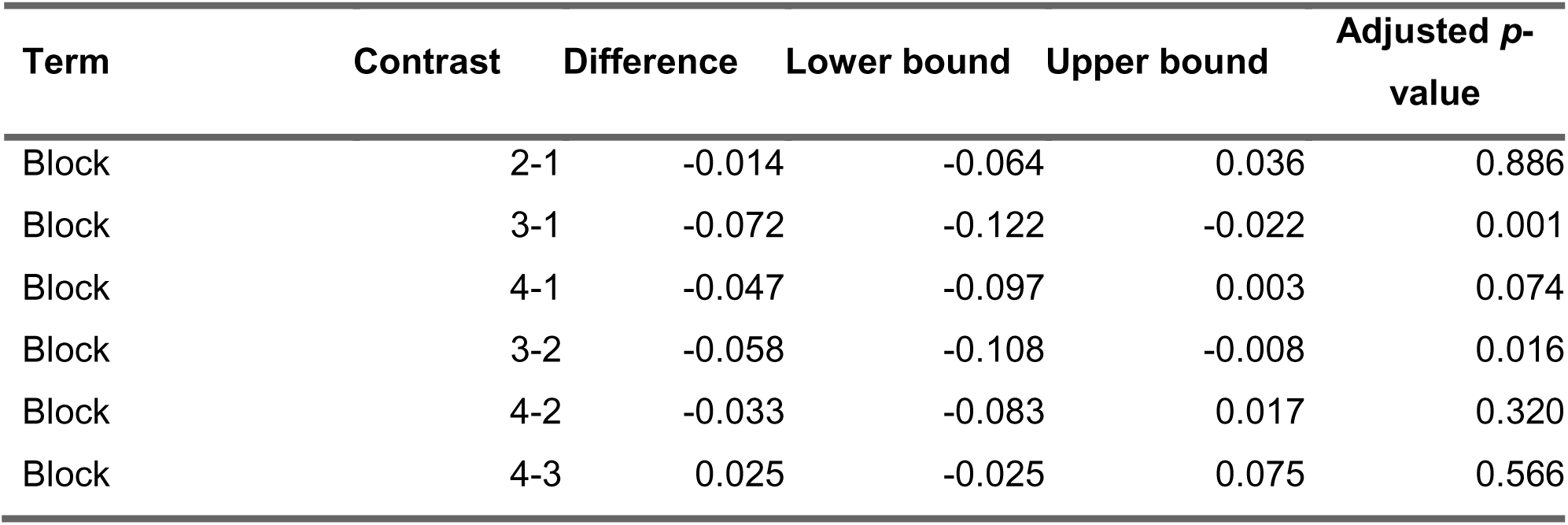
Tukey’s post-hoc analysis comparing blocks in the developmental lethal thermal limit data for *w*Au. Temperature values were transformed into categorical variables for this analysis.

**Figure S3.**
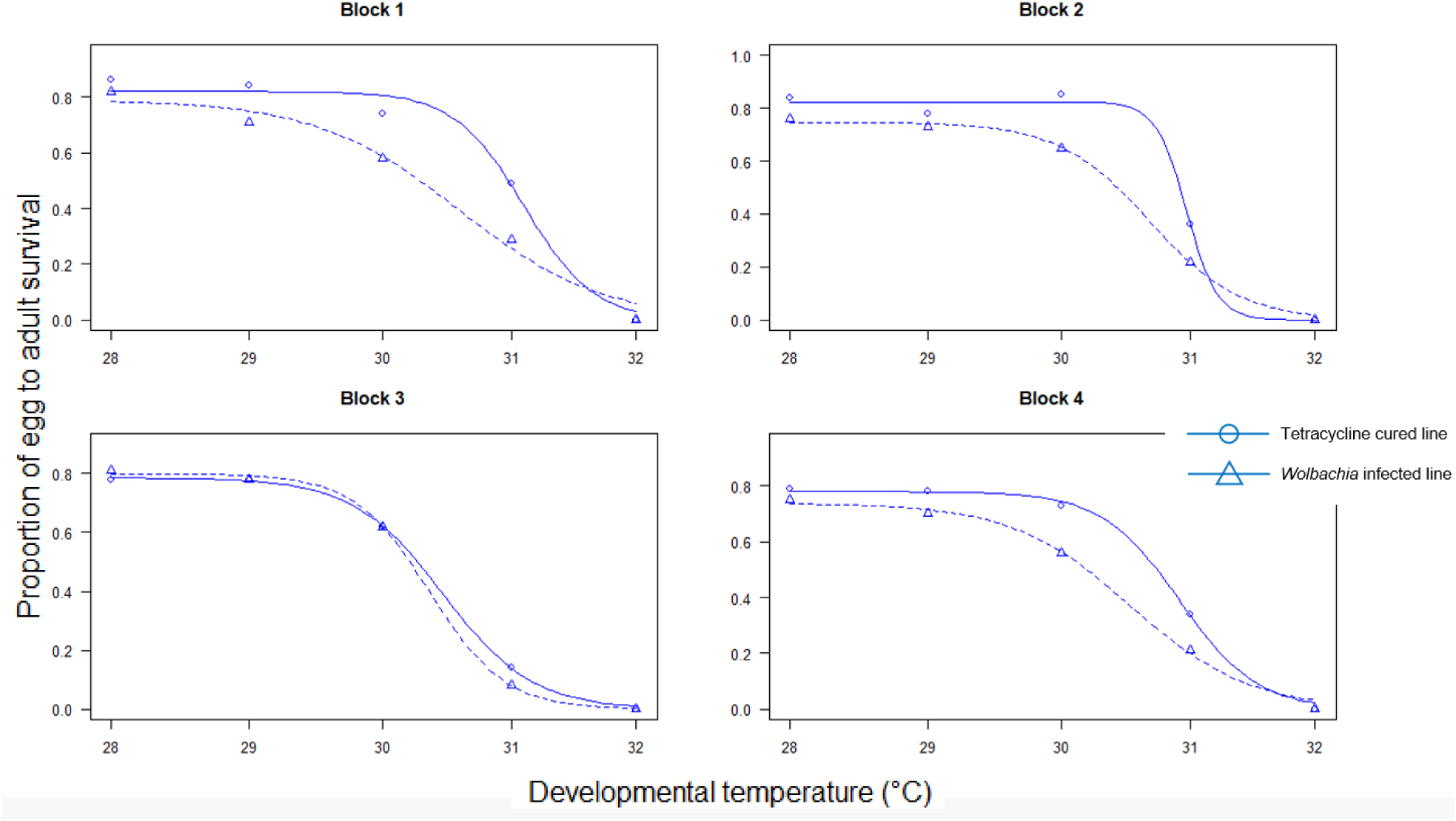
The proportion of surviving flies reared from egg to adult at different fluctuating developmental temperatures for the four blocks of the *w*Au line.

**Table S4.**
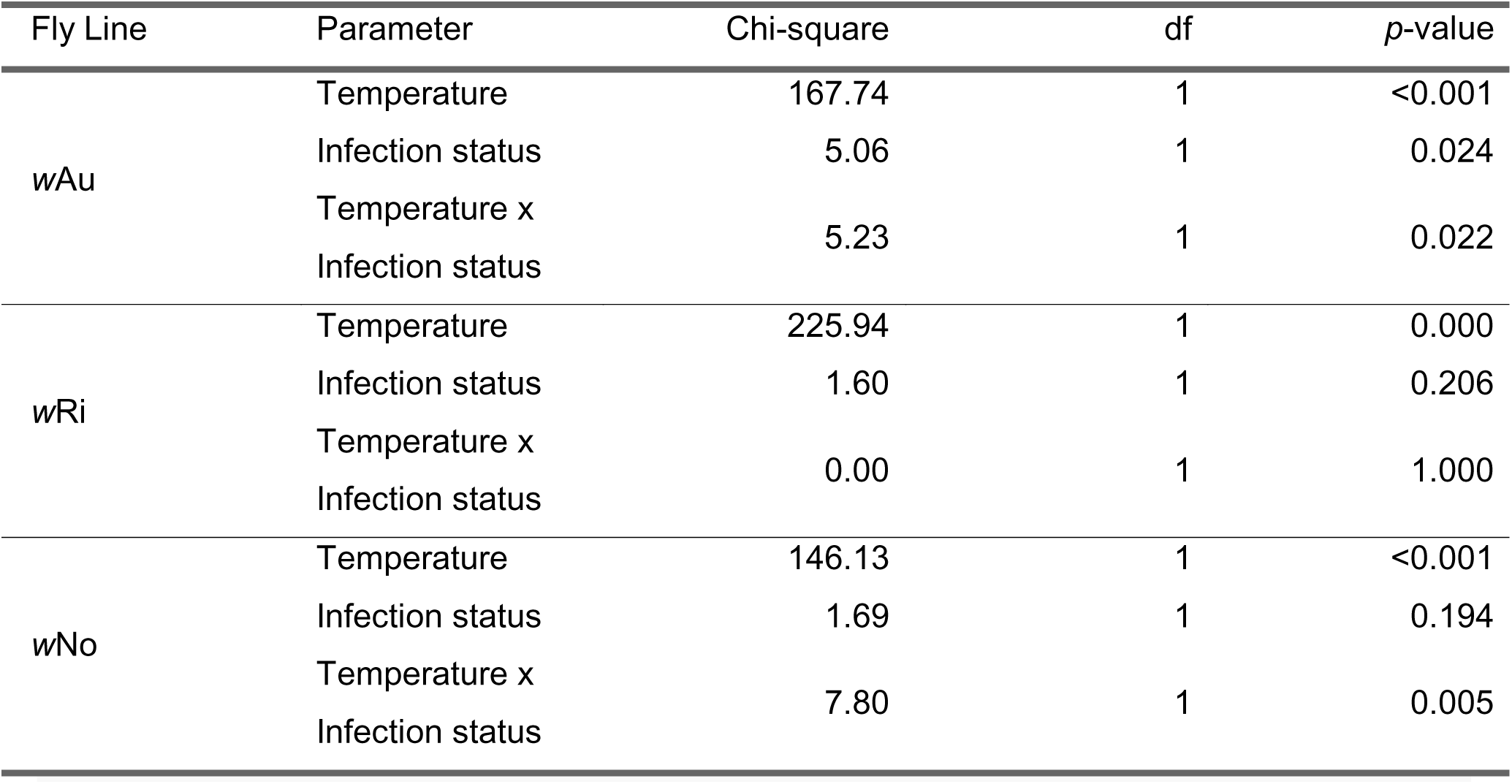
Two-way ANOVA assessing the impact of fluctuating developmental temperature, *Wolbachia* infection status and their interactions on male sterility (measured as the proportion of fertile males).

**Table S5.**
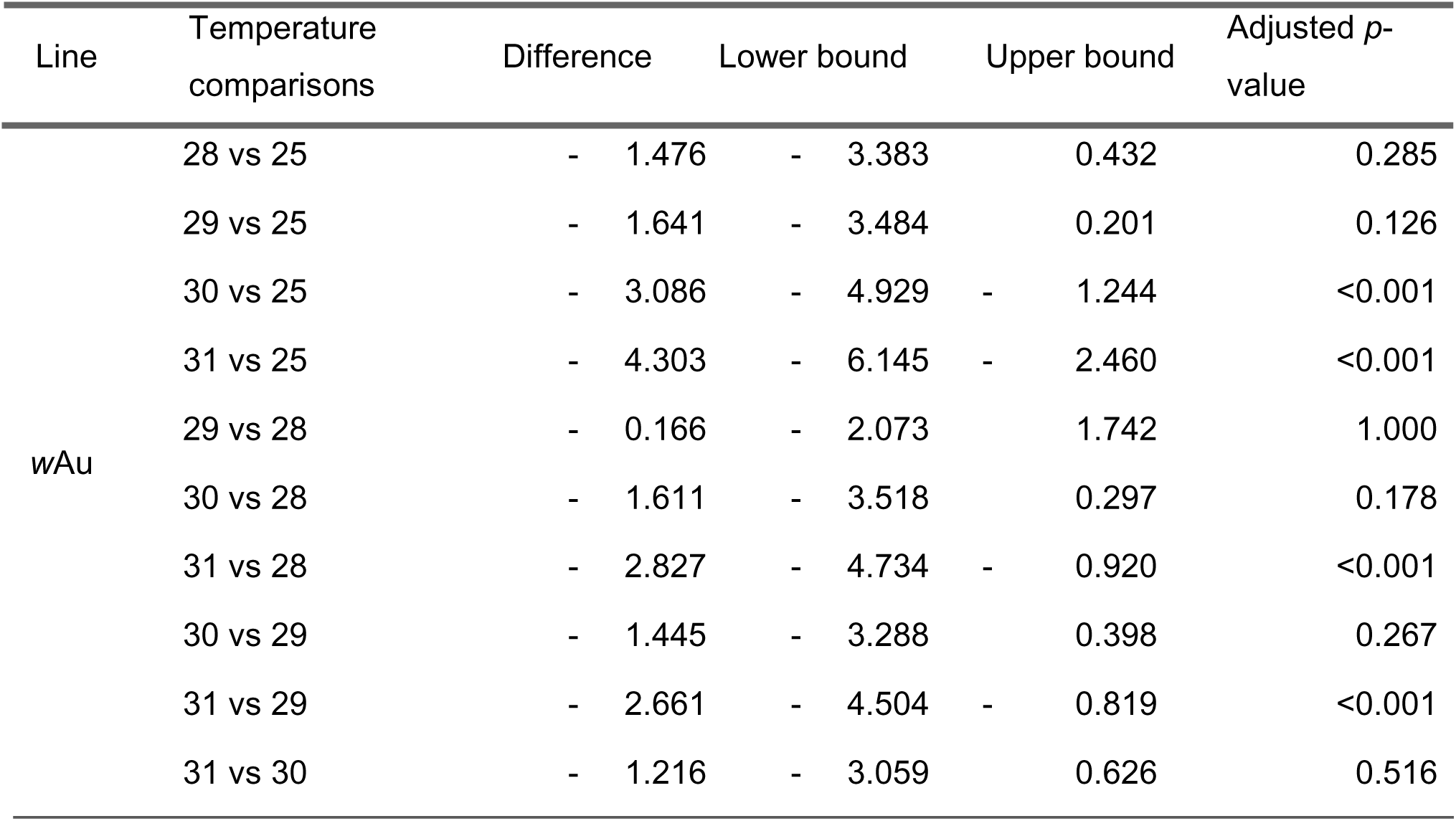
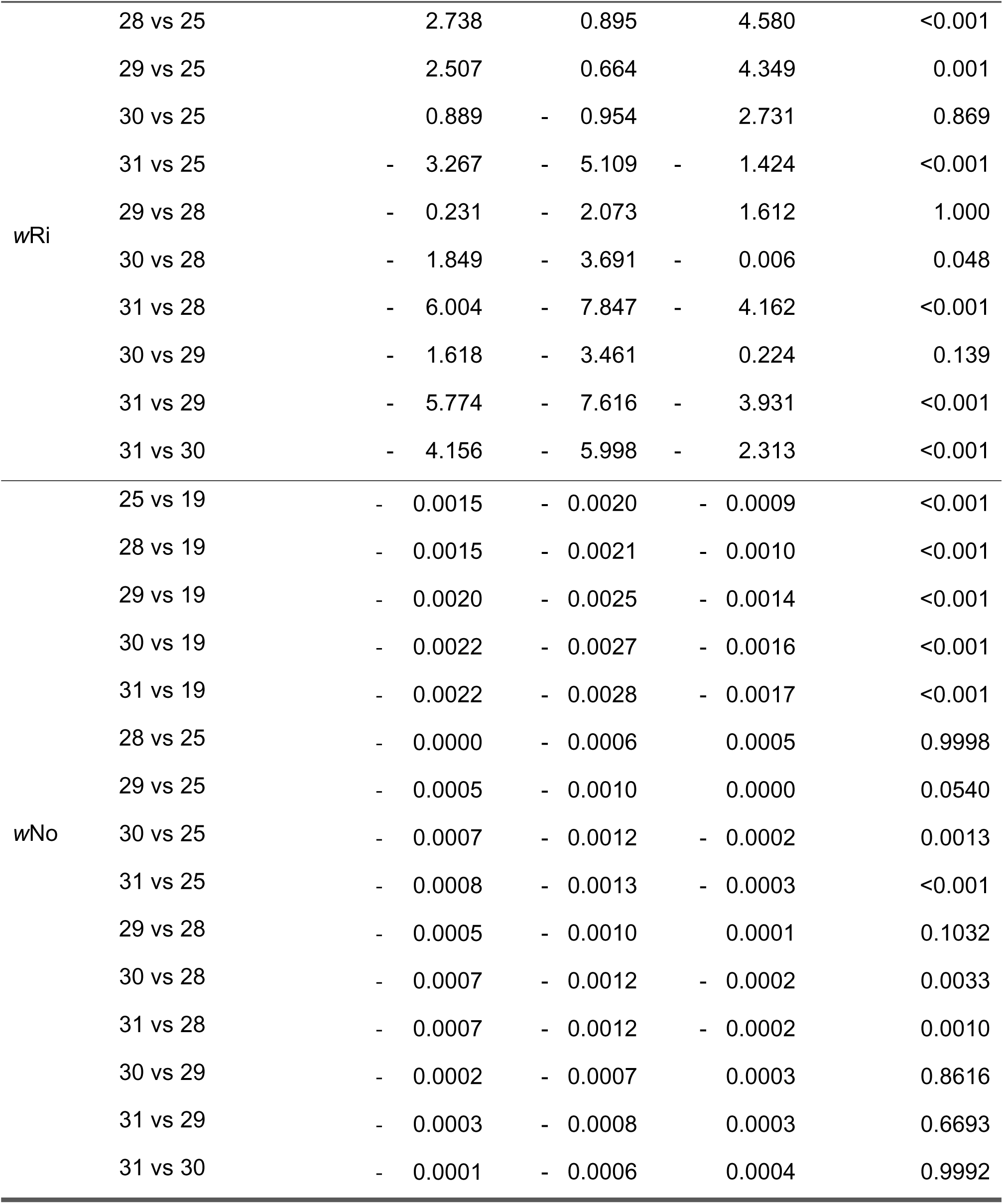
Tukey’s post-hoc analysis comparing *Wolbachia* densities between temperatures. Temperature values were transformed into categorical variables for this analysis.

| Line | Temperature comparisons | Difference | Lower bound | Upper bound | Adjusted p-value |
| --- | --- | --- | --- | --- | --- |
| wAu | 28 vs 25 | - 1.476 | - 3.383 | 0.432 | 0.285 |
|  | 29 vs 25 | - 1.641 | - 3.484 | 0.201 | 0.126 |
|  | 30 vs 25 | - 3.086 | - 4.929 | - 1.244 | <0.001 |
|  | 31 vs 25 | - 4.303 | - 6.145 | - 2.460 | <0.001 |
|  | 29 vs 28 | - 0.166 | - 2.073 | 1.742 | 1.000 |
|  | 30 vs 28 | - 1.611 | - 3.518 | 0.297 | 0.178 |
|  | 31 vs 28 | - 2.827 | - 4.734 | - 0.920 | <0.001 |
|  | 30 vs 29 | - 1.445 | - 3.288 | 0.398 | 0.267 |
|  | 31 vs 29 | - 2.661 | - 4.504 | - 0.819 | <0.001 |
|  | 31 vs 30 | - 1.216 | - 3.059 | 0.626 | 0.516 |

|  |  |  |  |  |  |  |  |  |  |
| --- | --- | --- | --- | --- | --- | --- | --- | --- | --- |
| wRi | 28 vs 25 |  | 2.738 |  | 0.895 |  | 4.580 |  | <0.001 |
|  | 29 vs 25 |  | 2.507 |  | 0.664 |  | 4.349 |  | 0.001 |
|  | 30 vs 25 |  | 0.889 | - | 0.954 |  | 2.731 |  | 0.869 |
|  | 31 vs 25 | - | 3.267 | - | 5.109 | - | 1.424 |  | <0.001 |
|  | 29 vs 28 | - | 0.231 | - | 2.073 |  | 1.612 |  | 1.000 |
|  | 30 vs 28 | - | 1.849 | - | 3.691 | - | 0.006 |  | 0.048 |
|  | 31 vs 28 | - | 6.004 | - | 7.847 | - | 4.162 |  | <0.001 |
|  | 30 vs 29 | - | 1.618 | - | 3.461 |  | 0.224 |  | 0.139 |
|  | 31 vs 29 | - | 5.774 | - | 7.616 | - | 3.931 |  | <0.001 |
|  | 31 vs 30 | - | 4.156 | - | 5.998 | - | 2.313 |  | <0.001 |
| wNo | 25 vs 19 | - | 0.0015 | - | 0.0020 | - | 0.0009 |  | <0.001 |
|  | 28 vs 19 | - | 0.0015 | - | 0.0021 | - | 0.0010 |  | <0.001 |
|  | 29 vs 19 | - | 0.0020 | - | 0.0025 | - | 0.0014 |  | <0.001 |
|  | 30 vs 19 | - | 0.0022 | - | 0.0027 | - | 0.0016 |  | <0.001 |
|  | 31 vs 19 | - | 0.0022 | - | 0.0028 | - | 0.0017 |  | <0.001 |
|  | 28 vs 25 | - | 0.0000 | - | 0.0006 |  | 0.0005 |  | 0.9998 |
|  | 29 vs 25 | - | 0.0005 | - | 0.0010 |  | 0.0000 |  | 0.0540 |
|  | 30 vs 25 | - | 0.0007 | - | 0.0012 | - | 0.0002 |  | 0.0013 |
|  | 31 vs 25 | - | 0.0008 | - | 0.0013 | - | 0.0003 |  | <0.001 |
|  | 29 vs 28 | - | 0.0005 | - | 0.0010 |  | 0.0001 |  | 0.1032 |
|  | 30 vs 28 | - | 0.0007 | - | 0.0012 | - | 0.0002 |  | 0.0033 |
|  | 31 vs 28 | - | 0.0007 | - | 0.0012 | - | 0.0002 |  | 0.0010 |
|  | 30 vs 29 | - | 0.0002 | - | 0.0007 |  | 0.0003 |  | 0.8616 |
|  | 31 vs 29 | - | 0.0003 | - | 0.0008 |  | 0.0003 |  | 0.6693 |
|  | 31 vs 30 | - | 0.0001 | - | 0.0006 |  | 0.0004 |  | 0.9992 |

**Table S6.**
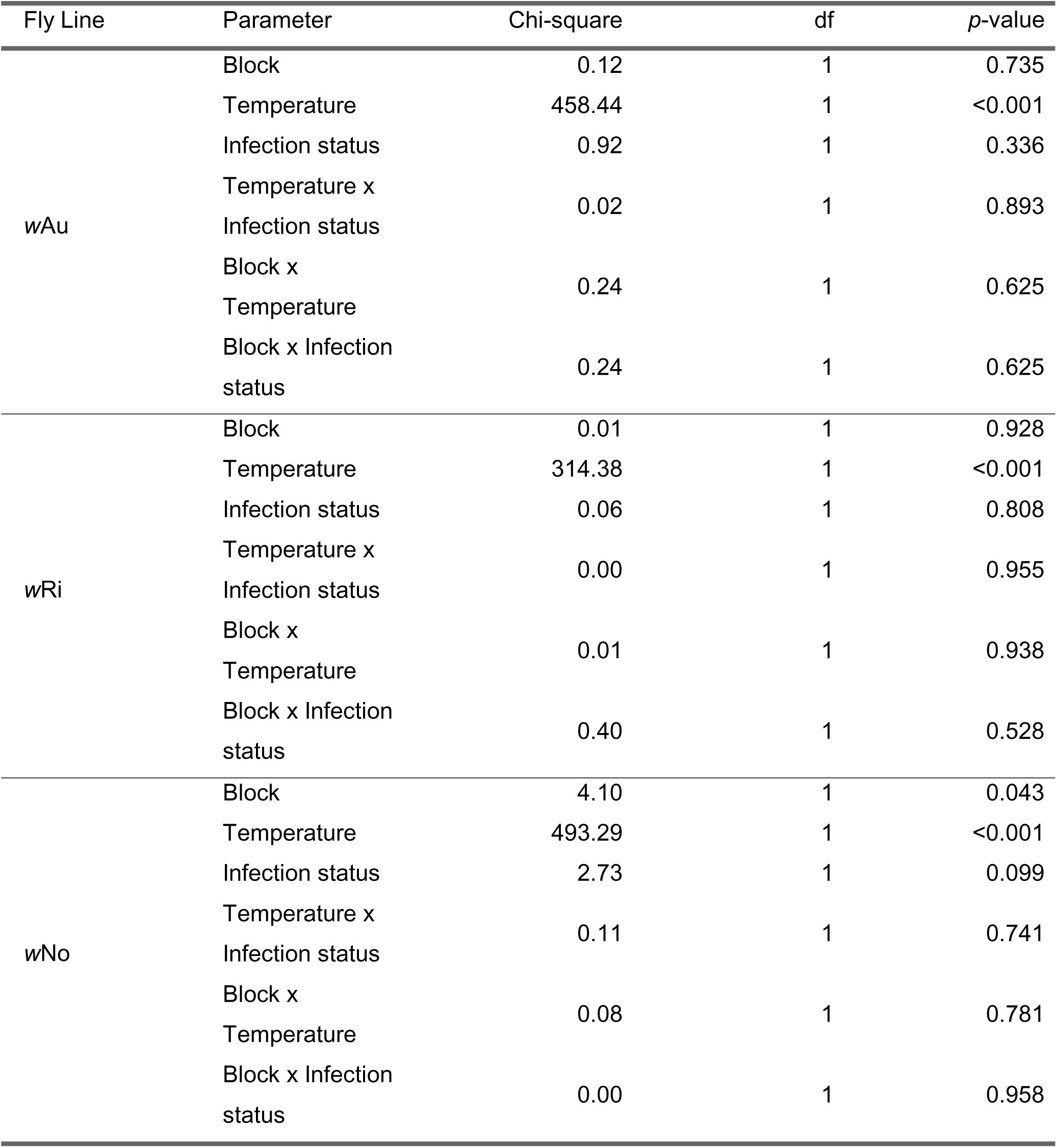
Two-way ANOVA assessing the impact of experimental block, heat shock temperature, *Wolbachia* infection status, and their interactions on the proportion of surviving male flies.

**Figure S4.**
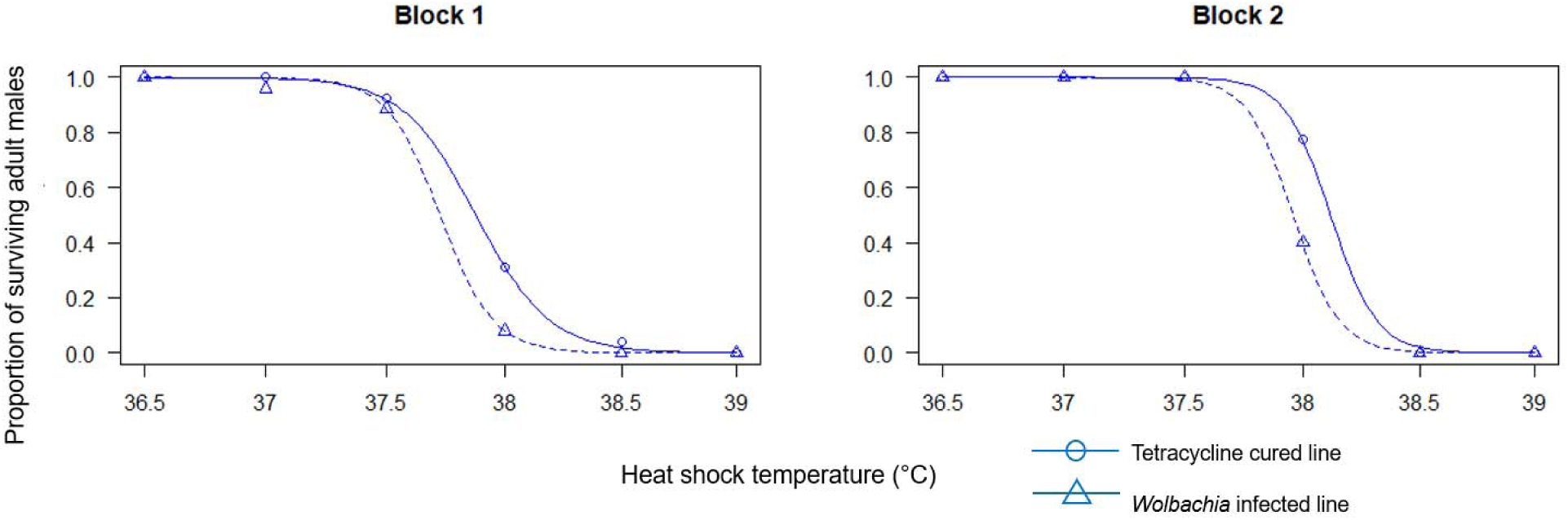
The proportion of surviving adult male flies exposed to a static 1-hour heat shock for the two blocks of the *w*No line.

**Table S7.**
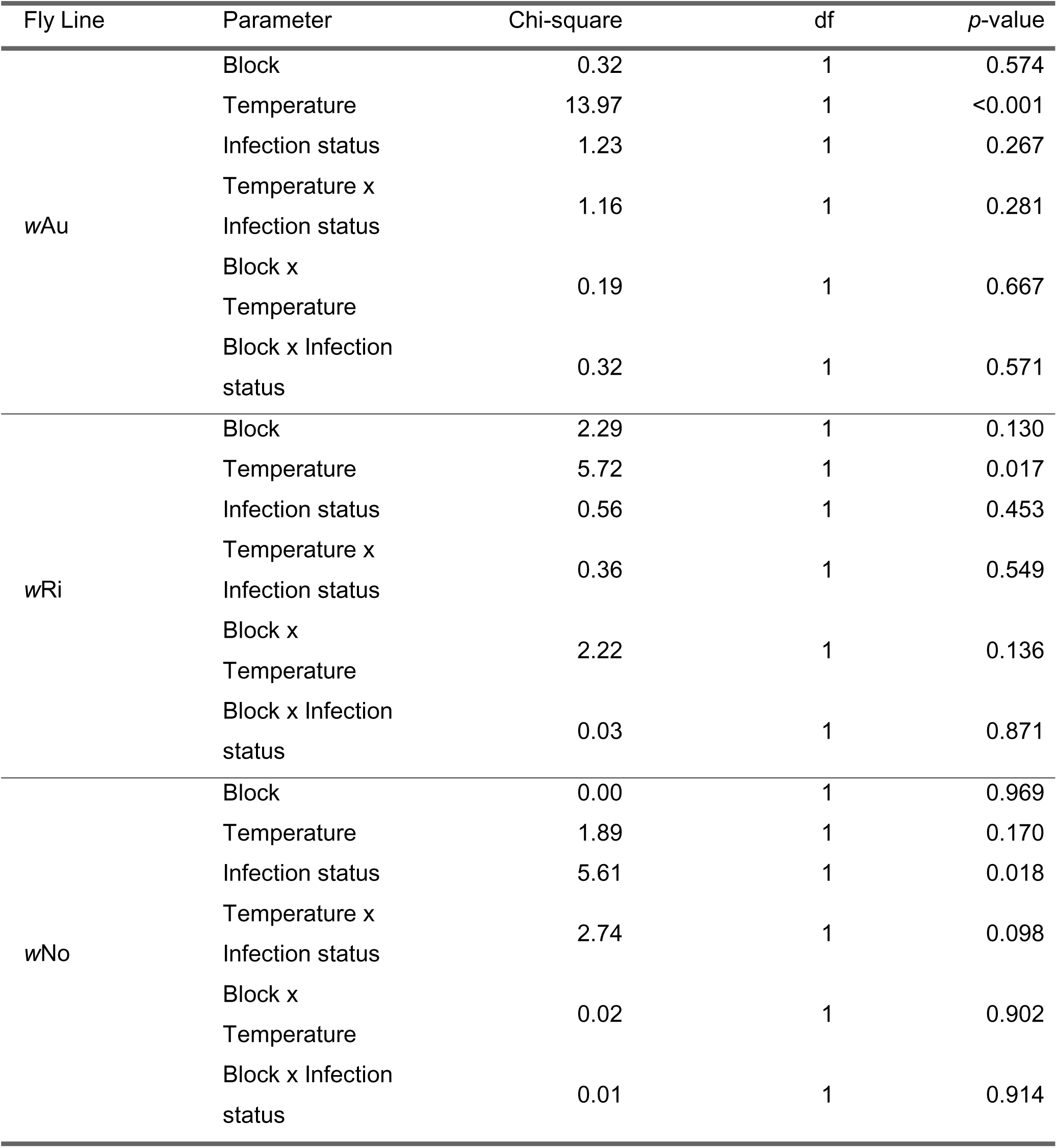
Two-way ANOVA assessing the impact of experimental block, heat shock temperature, *Wolbachia* infection status, and their interactions on adult male sterility (measured as the proportion of fertile males).

